# A genetic toolkit for the analysis of metabolic changes in *Drosophila* provides new insights into metabolic responses to stress and malignant transformation

**DOI:** 10.1101/711713

**Authors:** L Gándara, L Durrieu, C Behrensen, P Wappner

## Abstract

Regulation of the energetic metabolism occurs fundamentally at the cellular level, so analytical strategies must aim to attain single cell resolution to fully embrace its inherent complexity. We have developed methods to utilize a toolset of metabolic FRET sensors for assessing lactate, pyruvate and 2-oxoglutarate levels of *Drosophila* tissues *in vivo* by imaging techniques. We show here how the energetic metabolism is altered by hypoxia: While larval tissues that contribute directly to adult organs respond to low oxygen levels by executing a metabolic switch towards lactic fermentation, polytene tissues that are degraded during metamorphosis do not alter their energetic metabolism. Analysis of tumor metabolism revealed that depending on the genetic background, some tumors undergo a lactogenic switch typical of the Warburg effect, while other tumors don’t. This toolset allows for developmental and physiologic studies in genetically manipulated *Drosophila* individuals *in vivo*.

## Introduction

Carbohydrate catabolism is at the core of cellular bioenergetics (*1*). Pyruvate originated from glycolysis can be either reduced to lactic acid, or enter the mitochondria, where it is further oxidized to CO_2_ through the Krebs cycle reactions, providing reduced co-factors such as NADH or FADH_2_ that feed the electron transport chain, which generates the driving force for ATP synthesis *via* oxidative phosphorylation (OXPHOS) (*1*). Cells need to balance lactic fermentation and OXPHOS to cope with energetic and anabolic requirements upon changes in the environment. For example, mitochondrial OXPHOS becomes largely suppressed in hypoxia, as has been described in many models (*2–4*). To cope with this altered cellular physiology, many cells are capable of decoupling carbohydrate catabolism from mitochondrial OXPHOS by reducing pyruvate to lactate (Fig 1A). This metabolic status can be achieved through the regulation of a few enzymes or transporters that, acting together, control the metabolic flux. The main enzymes involved in this rewiring are *lactate dehydrogenase* (LDH), which converts pyruvate into lactate (*1*), and *pyruvate dehydrogenase kinase* (PDHK), which prevents pyruvate conversion into acetyl-CoA through the inhibition of the Pyruvate Dehydrogenase complex (*1*). Both enzymes, LDH and PDHK, are transcriptionally upregulated in hypoxia (*4, 5*). Likewise, other environmental challenges, such as nutrient deprivation (*1*) or osmotic shock (*6*), can also alter the metabolic profile of the cell.

**Figure 1:**
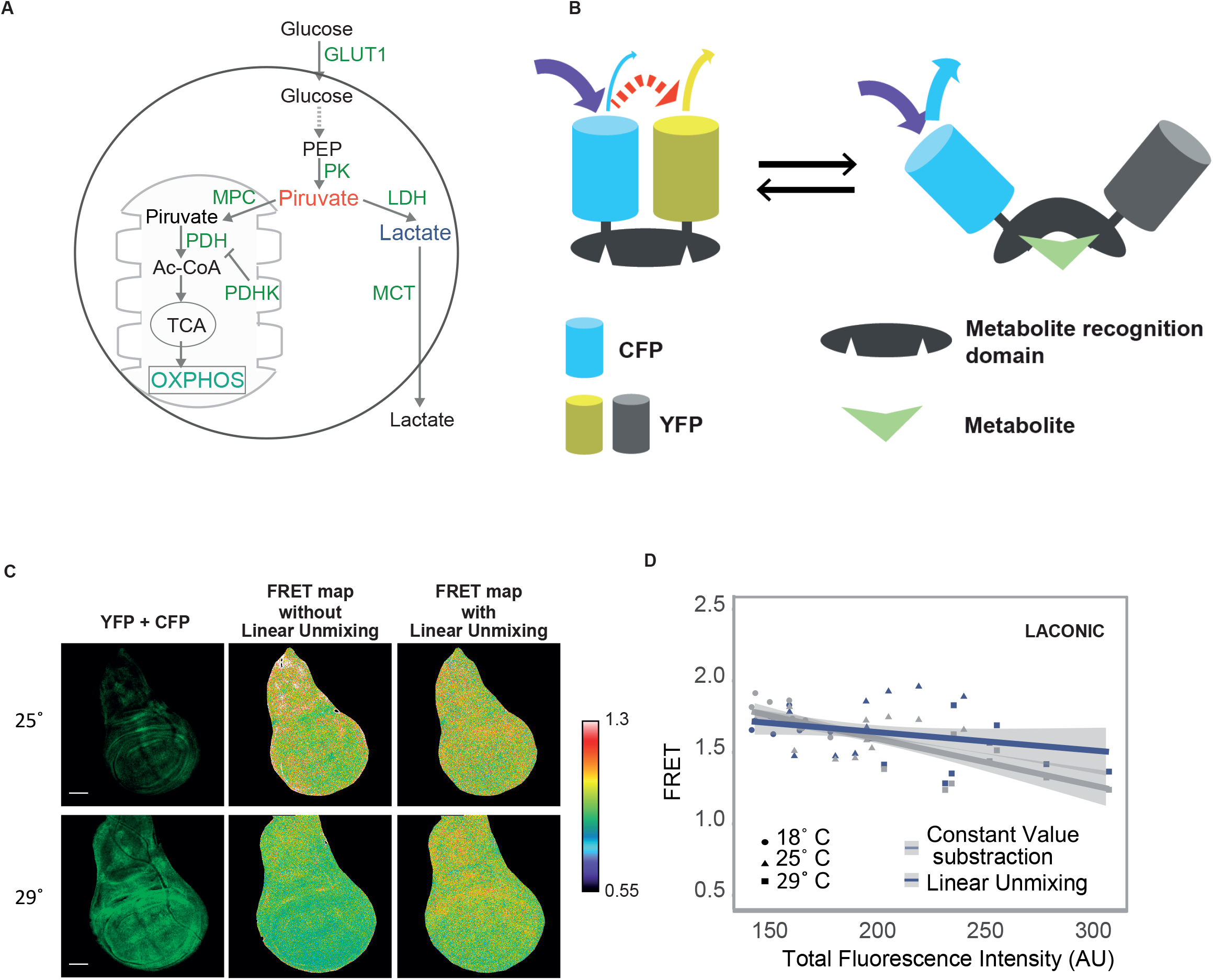
Sensors for studying cellular bioenergetics. A) Diagram of glucose catabolism. Glucose is broken down at the glycolysis and then can be fully oxidized to CO_2_ at the mitochondria (left), or alternatively, partially oxidized to lactate (right); pyruvate (red) stands as the branching point between the two alternative pathways. B) Schematic representation of the FRET sensors *(10-12)*. The donor and acceptor fluorophores, CFP and YFP, are represented in blue and yellow, respectively. Binding of the corresponding metabolite (green) to its binding domain separates CFP from YFP, and FRET ceases. C) FRET maps of the Laconic signal of wing imaginal discs in which different expression levels of the sensor were attained by using a tub-Gal4 driver at 25º C or 29º C. Note that the FRET signal obtained without linear unmixing is dependent on the expression levels of the sensor, while this dependency is largely suppressed after applying the linear unmixing algorithm. A high FRET signal in the color code shown on the right is associated to lower lactate levels. Scale bar: 50 μm. D) When a constant autofluorescence value was substracted throughout the whole image, the apparent FRET map (artefactual) depends on the expression levels of the sensor (grey curve; p= 1.07e-05). However, when the linear unmixing algorithm was utilized, the FRET map is no longer dependent on the expression levels of the sensor (blue curve; p= 0.114)

Several analytical methodologies are currently utilized to study metabolism in cell culture or animal tissues, provided sufficient amounts of material are available for preparing homogenates to perform biochemical analyses. These techniques encompass colorimetric assays to monitor enzymatic activity (*7*), chromatography (either GC, HPLC or UPLC) followed by mass spectrometry, or nuclear magnetic resonance to measure concentrations of different metabolites (*8*), as well as devices to study mitochondrial activity by assessing oxygen consumption rates (*9*). However, none of these widely used methodologies can monitor metabolic parameters in an intact organism with spatial resolution. The recent development of genetically-encoded fluorescent metabolic sensors opens this possibility, but imaging and data processing methods need to be improved to obtain reliable results in whole organisms or tissues.

Three FRET sensors that report levels of lactate (Laconic) (*10*), pyruvate (Pyronic) (*11*) or the Krebs cycle metabolite 2-oxoglutarate (2-OG) (*12*) have been developed in simple model systems such as bacteria or cell culture. In all three sensors, binding of the corresponding metabolite elicits a conformational change that separates the donor from the acceptor fluorophore, preventing resonant energy transfer (Fig 1B). Two of these sensors, Laconic and Pyronic, were then adapted to mice (*13*) and flies (*14*), although they have been utilized to monitor metabolic changes within single cells rather than to compare the metabolic status of a cell with that of the rest of the tissue (*10, 11*). We have independently generated transgenic fly lines for these three sensors, however when we tried to assess metabolite levels in individual cells in the context of intact organs, we faced artefactual results that precluded comparison between cells of endogenous metabolite levels. We present here an image processing method to obtain reliable FRET signals from the metabolic sensors Laconic, Pyronic and OGsor, opening the possibility to assess metabolite levels in developmental or physiologic studies in intact tissues or whole *Drosophila* organs. We employed these sensors to compare metabolic responses to hypoxia in different tissues, revealing that not all tissues undergo a lactogenic switch in the same manner. We also analyzed the occurrence of the Warburg effect in different experimental tumors induced in *Drosophila* larvae and found that this metabolic rewiring depends highly on their genetic background. We show that the tools and methods presented here can provide qualitatively distinct information to the biochemical approaches widely used in the field.

## Results and Discussion

### Optimized analysis of FRET sensor signals in full organs

#### Image processing algorithm to deal with autofluorescence

We generated fly lines in which the sensors Laconic, Pyronic or OGsor are expressed under control of UAS sequences, as well as a line that expresses Laconic ubiquitously under control of a Tubulin promoter (Methods). Initial attempts to employ these tools in whole *Drosophila* organs were unsuccessful due to diverse imaging artefacts: Most notably, the Laconic FRET signal seemed to correlate with expression levels of the sensor (Fig 1C). Induction of increasing levels of Laconic expression in 3^rd^ instar larval wing imaginal discs brought about an apparent FRET signal that tightly correlated with expression levels of the sensor (Fig 1C and D). In reporters such as these sensors, in which the donor/acceptor pair is part of the same protein (intramolecular FRET), the FRET signal should not depend on the sensor concentration in the sample, but only on their bound-to-unbound average ratio (*15*). This artefactual dependency on sensor expression levels prevents the use of the sensors in studies where different cells or tissues need to be compared, as expression levels of transgenes (the FRET sensors in this case) are never totally homogenous. As a possible cause of this artefactual behavior, we noticed that the background fluorescence varied with the temperature, and furthermore, that *Drosophila* larval tissues display high degree of pixel-to-pixel autofluorescence heterogeneity. Thus, correction of the background by subtraction of an average autofluorescence constant value, while effective in cell culture, is inappropriate *in vivo*.

To cope with autofluorescence heterogeneity, we adapted a *linear unmixing algorithm* (16), which estimates the autofluorescence contribution to the signal with single-pixel resolution (*supplementary information*). Briefly, the problem stems from the impossibility to estimate this heterogenic autofluorescence contribution to each pixel from an image of either the YFP or CFP channel. The linear unmixing algorithm allows dissecting these fluorescence sources, using a dedicated third image acquired in an emission window where only autofluorescence can be detected. After applying the linear-unmixing algorithm, the Laconic FRET signal does not depend any longer on sensor expression levels (Figure 1C and D).

In this manner, calculation of the FRET signal for each pixel defines a FRET map of a given 3^rd^ instar larval organ (Figure 2A). Analysis of different *Drosophila* larval tissues with this method revealed variations of lactate concentrations amongst individual cells (Figure 2B). Thus, by utilizing these sensors with a linear unmixing algorithm, the metabolic status of individual cells in the context of a whole *Drosophila* organ can be assessed.

**Figure 2:**
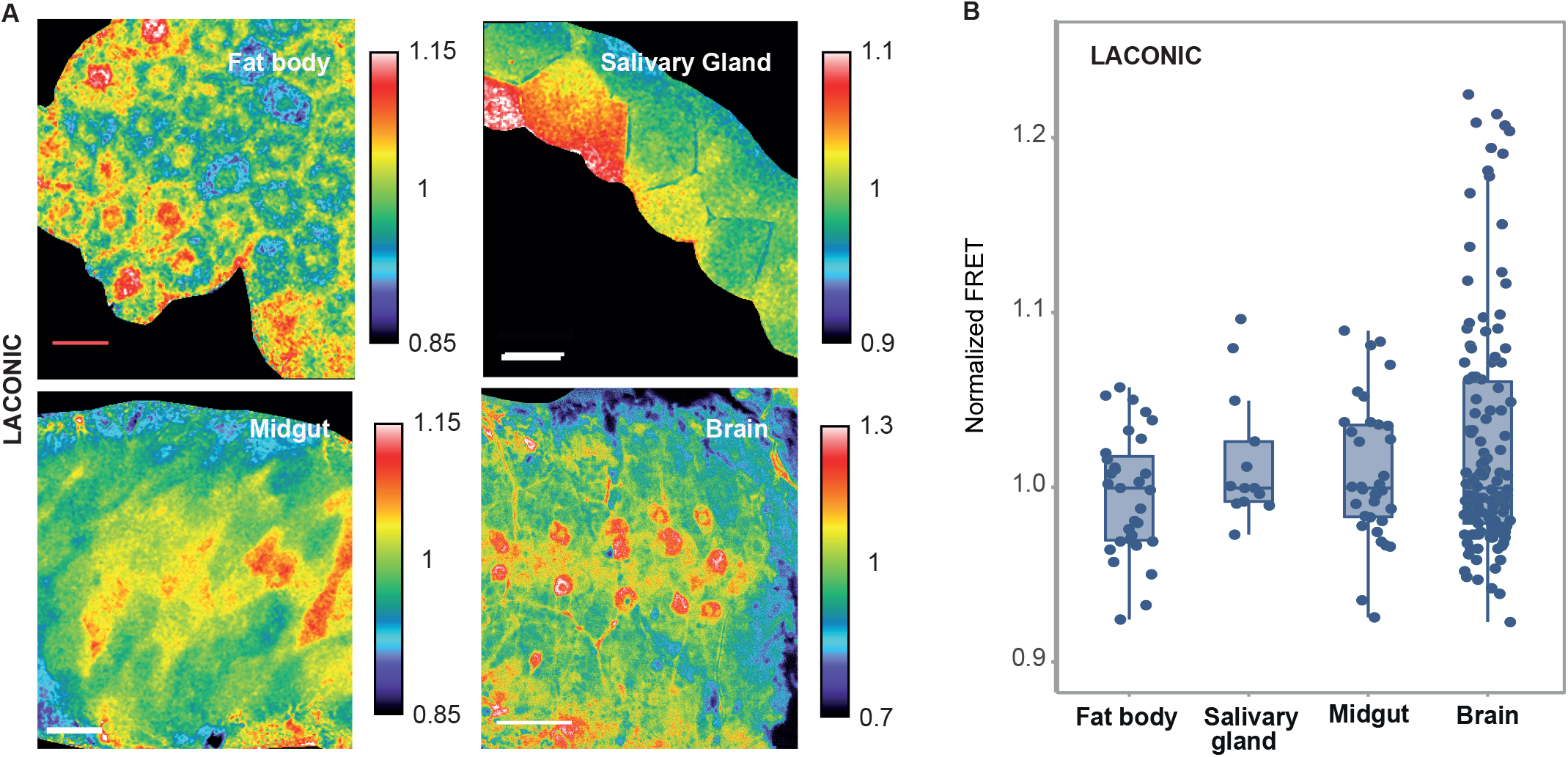
Laconic reveals single-cell variations of stationary lactate levels. A) Laconic FRET maps of a fat body, salivary gland, midgut and brain. In all four organs, cells with various different endogenous levels of lactate can be distinguished. Scale bar: 50 μm. B) The points represent the average FRET signal of each cell shown at the images of panel A). Data distribution is represented by the box and whiskers graph.

#### Validation and characterization of the FRET signal

We confirmed that energy transfer from the donor to the acceptor does indeed occur in our experimental setting. An assumption in FRET experiments is that the donor excitation wavelength does not induce direct excitation of the acceptor, and thus that all the fluorescence emission of the acceptor originates from energy transferred from the donor (Fig 1B). We therefore utilized a transgenic line that expresses only YFP (the acceptor) to rule out that the excitation wavelength of the donor (458 nm) can induce direct excitation of the acceptor in this experimental setting. As expected, this was not the case, implying that the YFP signal detected in flies bearing the sensor does indeed arise from FRET (Supplementary figure 1C).

As an additional control that FRET happens in transgenic flies carrying the sensors, we performed a set of *acceptor photobleaching* assays. With each of the sensors, we photoinactivated the acceptor fluorophore (YFP) by irradiating at high fluence a defined area of a larval wing imaginal disc with a wavelength at which the donor (CFP) is not stimulated (488 nm). If FRET does indeed occur, an increased emission of the donor is expected from the photobleached area upon irradiation at its excitation wavelength (458 nm). Comparison of the Laconic emission spectra of the irradiated region before and after YFP bleaching revealed an increase in CFP fluorescence (Sup Fig 1D and 1E), confirming that resonant energy transference effectively occurs between donor and acceptor. For Pyronic, lower fluence doses were employed to prevent undesired CFP photoinactivation, and in these conditions an increment of CFP fluorescence after photobleaching could be measured, which validates the Pyronic FRET signal (Sup Fig 1F). The YFP photobleaching assay on OGsor-expressing tissues only led to increased CFP emission in non-fixed material (Sup Fig 1G), so the experiments involving OGsor were carried out in live organs.

Next, to define the concentration range at which each of the sensors responds to its corresponding metabolite, we performed *ex-vivo* experiments. Wing discs, brains, fat bodies or salivary glands dissected from Laconic or Pyronic-expressing larvae were incubated in PBS buffer with increasing concentrations of lactate or pyruvate encompassing the expected physiological range, which rarely exceeds 10 mM (*17, 18*). Both sensors reported a reduction of the FRET signal proportional to the concentration of the corresponding metabolite (Fig 3A-E). Since 2-OG does not diffuse across the plasma membrane, to test OGsor responses we utilized the membrane permeable analog dimethyl-2-oxoglutarate (DM-2-OG). After reaching the cytosol, DM-2-OG is demethylated and converted into 2-OG, increasing its intracellular levels (*19*) and altering the OGsor FRET signal (Figure 3F).

**Figure 3:**
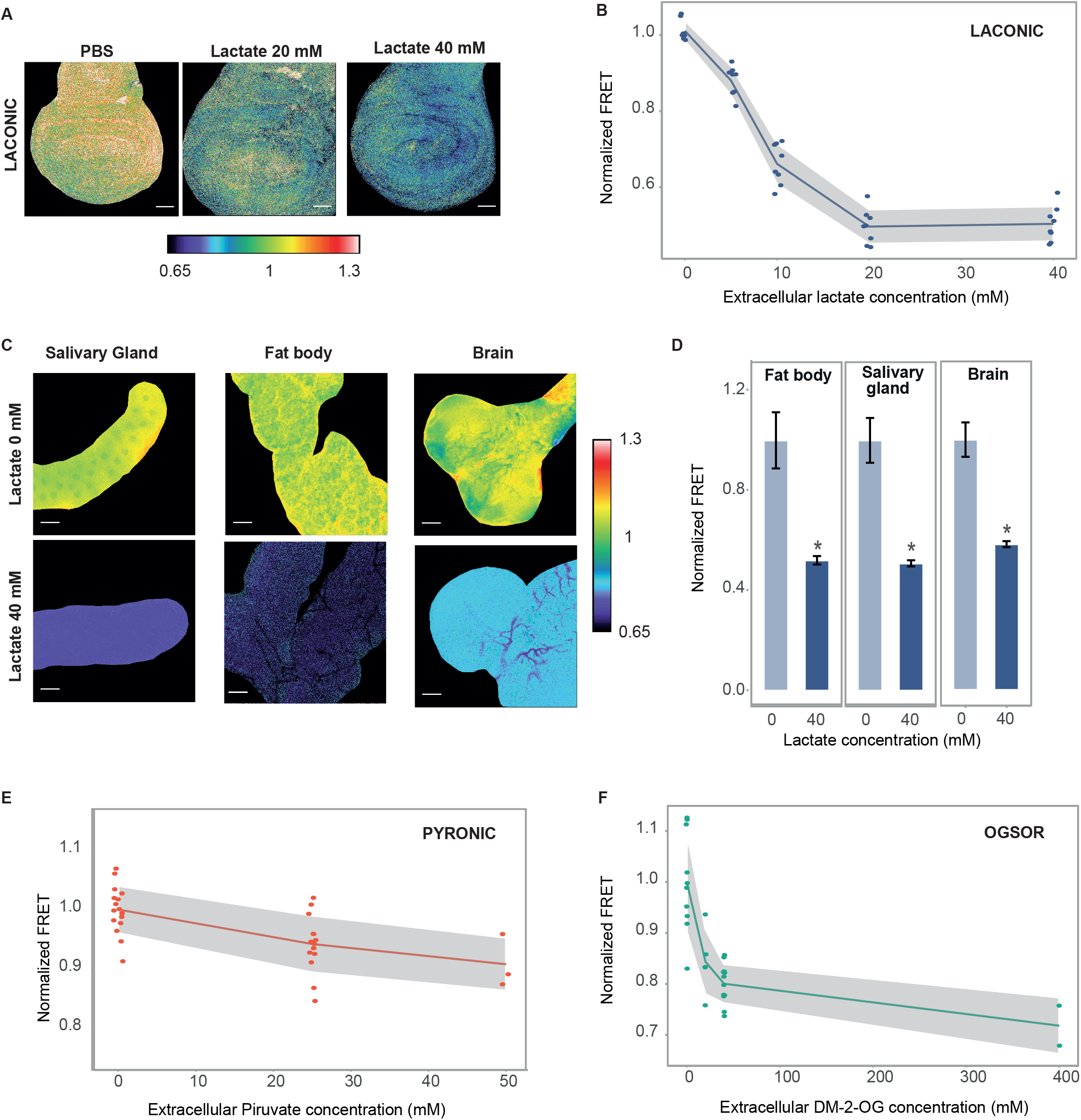
Response of the sensors to exogenously supplied metabolites. A) Laconic FRET maps of wing imaginal discs incubated or not in 20mM or 40mM lactate for 15 min. Scale bar: 50 μm. B) Quantification of the Laconic signal of the experiment of panel A. Each point represents the average value of a single imaginal disc. The grey area is limited by the SD of each data set. C) FRET maps of the indicated 3^rd^ instar larval organs with or without addition of exogenous 40 mM lactate. Scale bar: 50 μm. D) Quantification of the Laconic signal of the experiment shown in panel C). Data represent the media +/− SD; * p<0.05 Student’s T-test; n≥20 per group. E) Pyronic FRET signal from wing imaginal discs incubated or not with exogenous pyruvate for 15 min. Each point represents the average value of a single imaginal disc. The grey area is limited by the SD of each data set. F) OGsor FRET signal from wing imaginal discs incubated or not with exogenous dm-2-OG for 15 min. Each point represents the average value of a single imaginal disc. The grey area is limited by the SD of each data set.

### Environment-induced metabolic changes

Stress conditions are known to alter the cellular energetic metabolism (*1, 20, 21*), so we began by assessing the FRET signal of the three sensors in hypoxia. Wing discs of larvae exposed to hypoxic conditions for 16 h displayed increased lactate and decreased pyruvate levels, while we did not observe changes in the concentration of 2-OG (Fig 4A). These observations indicate that a lactogenic switch occurs in wing discs of hypoxic larvae. Interestingly, while the larval brain also executed a lactogenic switch, the salivary glands, midgut, and fat body did not (Fig 4B and C). Thus, a pattern seems to emerge, where non-polytene tissues -that will persist in the adult fly- become lactogenic in hypoxia, whereas polytene tissues, which will be degraded during metamorphosis, do not. The analyzed midgut region, which is composed of both non-polytene stem cells and polytene enterocytes (*22*), did make the lactogenic switch. Although further research is required to clarify this issue, it seems reasonable that genomic integrity needs to be especially preserved in tissues that will give rise to adult organs. Shut-off of mitochondrial OXPHOS, associated to a lactogenic switch in hypoxia, is expected to restrain accumulation of reactive oxygen species (ROS) (*23, 24*), thereby protecting cells from oxidative damage. Such a protective mechanism is probably not required in polytene tissues destined to histolysis during metamorphosis.

**Figure 4:**
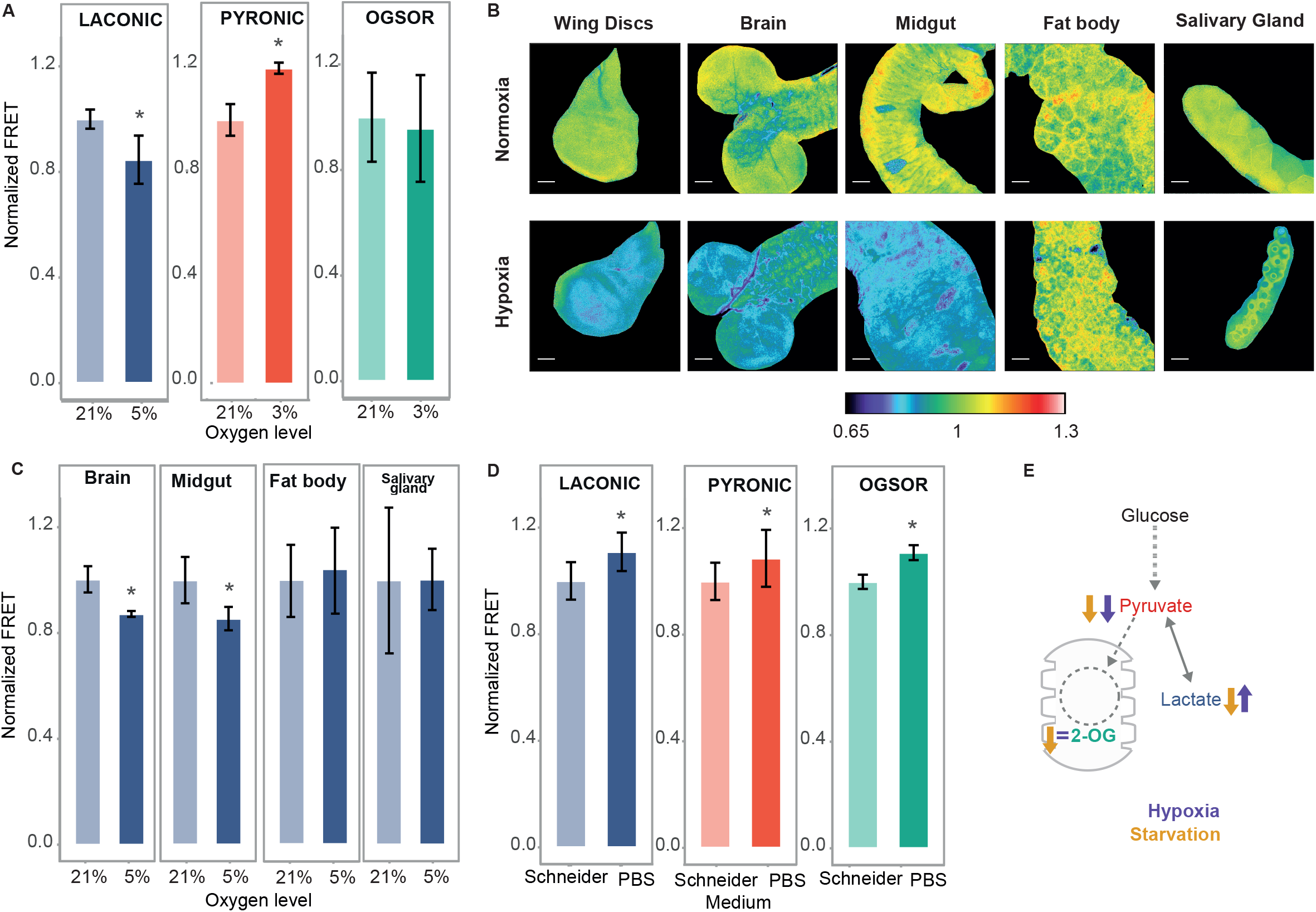
Metabolic rewiring upon O_2_ or nutrient deprivation. A) Laconic, Pyronic and OGsor FRET signal from wing imaginal discs from larvae exposed or not to hypoxia for 16 h before dissection and observation. Data represent the media +/− SD; * p<0.05 Student’s T-test. n≥20 per group. B) Laconic FRET maps of a wing disc, brain, midgut, fat body and salivary gland of 3^rd^ instar larvae exposed or not to hypoxia. Note that imaginal discs, the brain and midgut from larvae exposed to hypoxia increase their lactate levels, while salivary glands and the fat body do not. Scale bar: 50 μm. C) Quantification of the data of panel B); data represent the media +/− SD; * p<0.05 Student’s T-test; n≥20 per group. D) Laconic, Pyronic and OGsor FRET signal of 3^rd^ instar larvae wing imaginal discs incubated for 20 min in either Schneider medium or PBS prior to confocal analysis; data represent the media +/− SD; * p<0.05 Student’s T-test; n≥20 per group. E) Scheme of carbohydrate catabolism; variations of the metabolites monitored in this study upon hypoxia or starvation are indicated.

Nutrient deprivation is another environmental condition that can alter cellular bioenergetics. Starved *Drosophila* larvae experience a sharp reduction of catabolism of carbohydrates, proteins and lipids (*20*). This global metabolic constriction is thought to play a role in starvation resistance, albeit strong evidence supporting this hypothesis is still elusive (*25, 26*). The way in which starvation-induced catabolic inhibition affects the stationary levels of the metabolites analyzed here has not been explored at the cellular level, although lactate concentrations in the whole larva have been shown to diminish upon protein starvation (*27*). Larvae expressing each of the three sensors were subjected to 6 h starvation, a condition strong enough to induce autophagy (*28*). No alterations of the FRET signal from any of the sensors could be detected (Sup Fig 2.A). These results suggest that larval starvation does not impinge on intracellular levels of Lactate, Pyruvate or 2-OG, probably due to physiological compensation mechanisms capable of maintaining cellular homeostasis. The Laconic signal also remained unaltered upon a longer (18 h) starvation period in all of the analyzed organs (Sup Fig 2.B). Noteworthy, these tools cannot report on every kind of metabolic rewiring: only those cases that lead to altered steady-state levels of the monitored metabolites (lactate, pyruvate or 2- OG) are visible to the sensors, while compensated flow changes might still occur.

If systemic compensation mechanisms do indeed account for intracellular stability of Lactate, Pyruvate and 2-OG steady-state levels, variations of these metabolites might be observed in tissues subjected to starvation *ex vivo*. We incubated wing imaginal discs dissected from larvae in Schneider (rich) medium or in PBS (extreme starvation) for 15 minutes. Starved imaginal discs displayed decreased levels of lactate, pyruvate and 2-OG, as reported by each of the sensors (Fig 4D). Thus, an overall nutrient restriction results in reduction of all three metabolites, which probably reflects the decreased metabolic flux characteristic of the starvation response.

In summary, whilst hypoxia induces a lactogenic switch in non-polytene tissues that involves increased lactate and reduced pyruvate levels, extreme starvation leads to decreased concentrations of the three metabolites, lactate, pyruvate and 2-OG, reflecting an overall reduced metabolic flux (Fig 4E).

### Genetic manipulations of the bioenergetic metabolism

We analyzed the Laconic signal after manipulating levels of the glucose transporter Glut1, the Mitochondrial Pyruvate Carrier (MPC), and the Monocarboxylate Transporters, (MCTs) Silnoon and Chaski. We also manipulated the expression of key enzymes of the energetic metabolism such as pyruvate kinase (PK), pyruvate dehydrogenase kinase (PDHK) or lactate dehydrogenase (LDH) (Figure 1A). All the above genes are essential for regulation of the glycolysis/OXPHOS balance in diverse physiological or pathological contexts (Table 1). Neither silencing nor over-expression of any of these individual genes elicited alterations of the Laconic FRET signal in wing discs (Sup Fig 3). In an attempt to overcome metabolic robustness, and force alterations of intracellular lactate levels, we induced combinations of two simultaneous genetic manipulations expected to act synergistically. As depicted in Figure 5A, silencing of MPC combined with overexpression of PDHK or LDH led to increased intracellular lactate levels in the cells where those genes were altered (Fig 5A and B).

**Table 1:** Key metabolic enzymes or transporters whose transcriptional deregulation has been reported to alter the energetic metabolism. The table summarizes their physiological role in metabolism as well as the kind of alterations of these genes in tumorigenesis.

| Protein | Role in the energetic metabolism | Alterations reported in cancer contexts | References |
| --- | --- | --- | --- |
| GLUT1 | Glucose transport through plasma membrane | Activation of isoforms 1 and 3 in mammalian tumor models<br>Its silencing, concomitant with LDH, reduces the tumor phenotype observed upon Notch activation in *Drosophila* imaginal discs | (34, 39) |
| Piruvate Kinase (PK) | Conversion of Phosphoenolpyruvate into pyruvate | Increase in PKM2 isoform in several mammalian tumor models | (40, 41) |
| Mitochondrial Pyruvate Carrier (MPC) | Mitochondrial transporters of pyruvate | Inhibition of isoforms 1 and 2 in mammalian tumor models<br>Its silencing in *Drosophila* leads to larval lethality when development occurs in media that lacks carbon sources other than sucrose | (42, 43) |
| Monocarboxylate Transporter (MCT) / Silnoon and Chaski | Transporters of lactate and pyruvate in the plasma membrane | Heightened expression of isoform 4 in mammalian tumor models | (44) |
| Pyruvate Dehydrogenase Kinase (PDHK) | Inhibition of the Pyruvate Dehydrogenase complex | Increase in Isoform 1 in several mammalian tumor models | (45, 46) |
| Lactate Dehydrogenase (LDH) / Impl3 | Reduction of pyruvate and lactate synthesis | Increase in LDHA isoform in several mammalian tumor models<br>Transformation from hyperplasia to neoplasia depends on LDH activity | (37, 47) |

**Figure 5:**
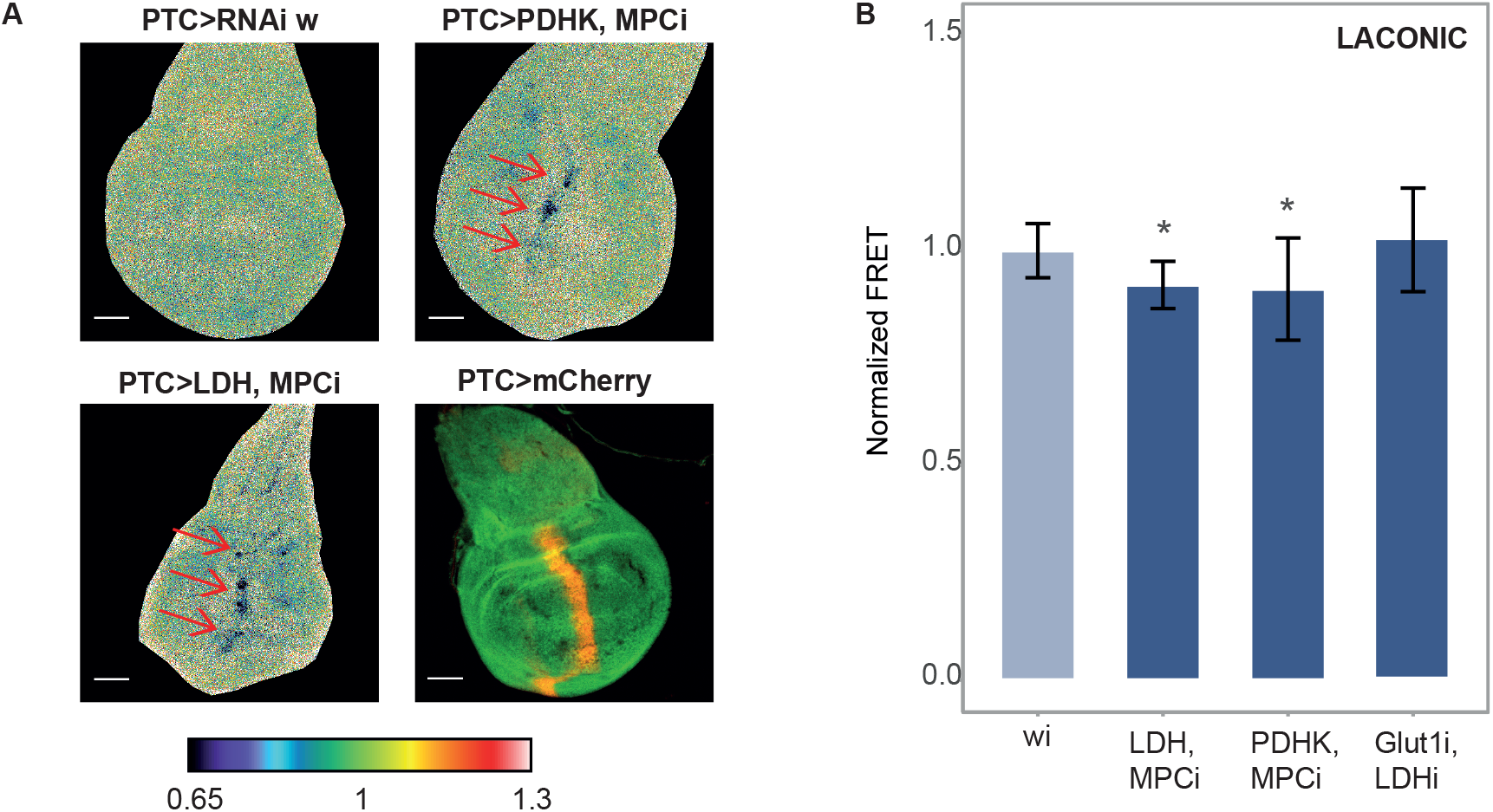
Tissue bioenergetic metabolism can be genetically rewired. A) Laconic FRET maps of 3^rd^ instar larvae wing discs, in which the expression of the indicated genes has been manipulated with Patched-Gal4. The arrows indicate cells of the disc in which genetic manipulations provoked an increase of lactate levels. The Patched expression domain is revealed by mCherry expression. B) Variation of the Laconic FRET signal from 3^rd^ instar larvae wing discs where the combined genetic manipulations indicated on each column were carried out at the posterior compartment of the disc with an En-Gal4 driver. Data represent the media +/− SD; * p<0.05 Dunnet’s Test; n≥20 per group.

The energetic metabolism is a paradigmatic case of biological robustness (*29, 30*), and thus it is not surprising that the metabolic outcome is not altered after manipulation of single metabolic genes. For example, when lactate synthesis rates are below a certain threshold, MCTs may act as a drain for lactate, avoiding intracellular accumulation. If this is the case, at lactate levels that are above that threshold, flux through MCTs could become saturated, provoking intracellular lactate accumulation. Thus, increased lactate levels that we observe by combining manipulations of the MPC and either LDH or PDHK might reflect this dynamics.

### The Warburg effect in *Drosophila*

Even under conditions of oxygen availability, human tumor cells undergo a metabolic switch towards glycolysis, known as the Warburg effect (*31*), and recent reports indicate that these metabolic alterations are recapitulated in *Drosophila* experimental tumors (*32, 33*). We tested whether Laconic can detect a Warburg-like metabolic switch in *Drosophila* experimental tumors generated by a variety of genetic strategies. Interestingly, and in line with earlier indirect observations (*33*), only some specific genetic manipulations elicited the Warburg effect.

Constitutive activation of the Notch pathway in wing imaginal discs produce deregulated growth and hyperplastic tumors (*34*), which depend on the induction of glycolysis-related genes such as LDH, Hexokinase A, and Glut1. Using Laconic, we analyzed intracellular lactate levels in these developing tumors, and found no significant differences in comparison to normal tissues (Fig 6A and B). In contrast, overexpression of an activated variant of the PDGF/VEGF receptor homolog PVR (*33*), or of an activated version of Ras, provoked a reduction of the Laconic FRET signal (Fig 6A and B), indicating that lactate accumulated intracellularly. These observations are consistent with a previous report (*33*), in which tumors induced through the same strategies displayed overexpression of LDH. We also employed Laconic to analyze tumor models in which metabolism had not been explored before. *Lethal (2) giant larvae* (l2gl) is a membrane-associated protein that regulates both cell proliferation and the epithelial polarity, and whose mutants have been shown to produce neoplastic tumors (*35*). Silencing of l2gl led to the formation of lactogenic tumors (Fig 6A and B).

**Figure 6:**
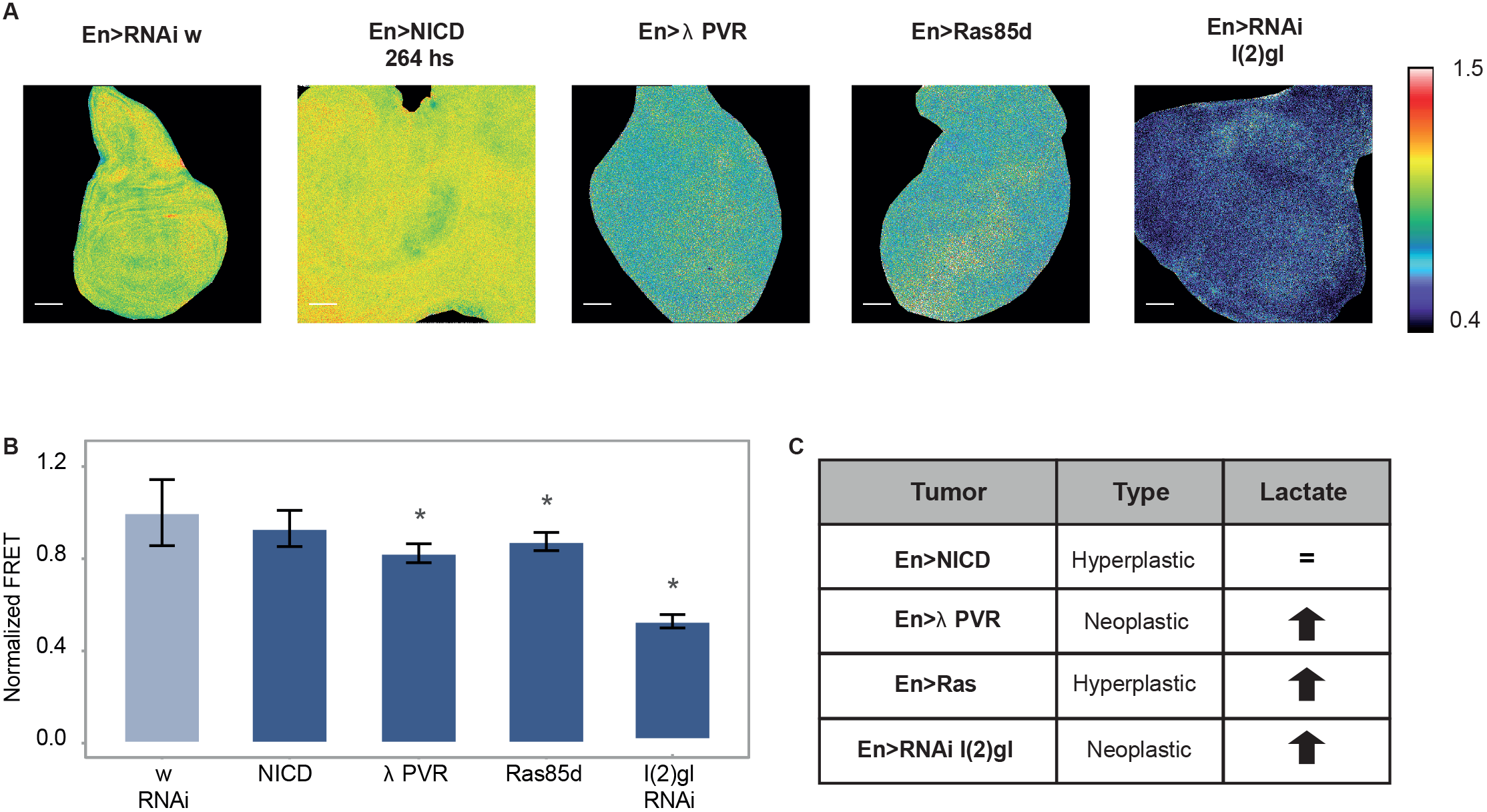
The Warburg effect in experimental tumors induced by different genetic strategies. A) Laconic FRET maps of tumors induced in imaginal wing discs by manipulating gene expression with an en-Gal4 driver. Scale bar: 50 μm. B) Quantification of the results of panel A); data represent the media +/− SD; * p<0.05 Student’s T-test; n≥20 per group. In A and B data from each tumor was normalized to the mean value of the w RNAi genotype analyzed in parallel. C) Summary of the lactate levels in each of the tumors.

Tumors in *Drosophila* are described as either *hyperplastic* or *neoplastic*. Whilst the former category is defined by deregulated proliferation with no alterations of cell shape or tissue polarity (*36*), neoplastic tumors encompass rounded cells that lost polarization and associate with an altered architecture of the tissue (*36*). It has recently been reported in tumors induced by overexpression of the EGF receptor (*37*), that a lactogenic switch is associated to transformation of hyperplastic to neoplastic tumors. In line with this finding, our observations that the neoplastic tumors induced by l(2)gl or activated PVR undergo a lactogenic switch suggest that this metabolic requirement might be, in fact, a general feature of neoplastic growth. However, the fact that Ras-induced tumors are also lactogenic indicates that the metabolic switch is not restricted to neoplasia. Taken together, our results indicate that tumors of different genetic origin display different metabolic properties (Fig 6C); a facet of tumor biology that is just starting to be explored.

### Final remarks

We have shown here that metabolic FRET sensors can be used to characterize the metabolic response of a single organ upon exposure to stress conditions. We found not only that wing imaginal discs respond in a different manner whether the stress is induced by nutrient or oxygen deprivation, but also that different organs react differently to hypoxia. We have also shown that Laconic is capable of reporting an altered lactate concentration produced by genetic manipulation of key metabolic enzymes in a restricted domain of wing discs. These results pave the way to a systematic exploration of the effect of single genes on cell metabolic status *in vivo*. Reverse genetics analyses utilizing these metabolic sensors could shed light on the role that each enzyme and transporter plays in metabolic responses to different physiologic or pathologic conditions.

## Methods

### Fly lines and stocks

The following fly stocks were obtained from the Bloomington *Drosophila* Stock Center (https://bdsc.indiana.edu/): UAS-NICD (52008), UAS-λPVR (58428), UAS-Ras85d (4847) tub‐Gal4 (5138), en-Gal4 (1973), ptc-Gal4 (2017), Glut1 RNAi (40904) l(2)gl RNAi (38989) and white RNAi (33613). The following stocks were from the Vienna *Drosophila* Research Center (https://stockcenter.vdrc.at/): LDH RNAi (110190), PDHK RNAi (37966), MPC RNAi (103829), PK RNAi (49533) Chk (MCT) RNAi (37141) and Silnoon (MCT) RNAi (106773). The UAS-LDH line was obtained from the Zurich ORFeome Project (https://flyorf.ch/).

The following lines were generated in this work: UAS-Laconic, UAS-Pyronic, UAS-OGsor, tub-Laconic, UAS-PDHK.

### Cloning and transgenic lines generation

Transgenic lines bearing UAS-Laconic, UAS-Pyronic, UAS-OGsor and tub-Laconic were generated by phiC31-mediated site-directed integration on the 58A landing site. UAS-PDHK, on the other hand, was integrated into the 86F landing site.

The ORF of Laconic and Pyronic were subcloned into the pUASt.attB vector using XhoI and XbaI. OGsor was subcloned in the same vector using BglII and NotI. Laconic was also subcloned into pss193, downstream of the tubulin promoter, using BamHI and XbaI. Finally, the ORF of PDHK was obtained from the Drosophila Genomics Resource Center (https://dgrc.bio.indiana.edu; #BS06809), and then subcloned into pUASt.attb using NotI and XbaI.

### Microscopy setup

Images of the three relevant channels were obtained simultaneously using the QUASAR detection unit of the Zeiss 710 or 880 microscopes. The emission windows that define these channels are: donor (CFP) channel, 490 +/− 5 nm; acceptor (YFP) channel, 530 +/− 5 nm; and autofluorescence (A) channel, 600 +/− 5 nm. Emission spectra, whenever required, were obtained using a Zeiss LSM510 Meta Confocal Microscope with monochromator. Sensors were excited at 405 nm or 458 nm.

### Starvation and Hypoxia treatment

For larval starvation, 3^rd^ instar larvae were collected from their regular medium, and placed in 2% agar plates supplemented with 3% sucrose, for 6 h as previously reported (*28*). For hypoxia treatments the proportions of oxygen and nitrogen were regulated in a Forma Scientific 3131 incubator. Third instar larvae were subjected to hypoxia for 16 h before dissection and observation under the confocal microscope. Larvae were dissected in PBS and then fixed in 4% formaldehyde (Sigma, St. Louis, MO, USA) for 120 min at room temperature. After washing three times for 10 minutes in PT (PBS, 0.3% Triton-X 100), the required organs were separated and mounted in Mowiol (Calbiochem, Merck & Co, New Jersey, USA).

### Statistical analysis

Data are expressed as mean ± standard deviation (SD). When comparing between two conditions, the Student’s T-test was employed. Normality was tested using the Shapiro–Wilks test. If data did not followed a normal distribution, the Mann-Whitney test was used instead. For multiple comparisons, one-way analysis of variance (ANOVA) followed by Dunnet’s test was performed; in this case, data were tested for normality with the Shapiro–Wilks test and variance homogeneity with the Levene test. Box-Cox transformations were employed whenever normality or homocedasticity requirements were not satisfied. A p<0.05 was considered statistically significant. Statistical analyses were executed using GraphPad Prism, version 5.03 (GraphPad Software).

## Acknowledgments

We thank Luis Felipe Barros and Bang-Ce Ye for reagents, Nicolás Frankel, Cecilia D’Alessio, Andrés Garelli, Valeria Levi, Hernán Grecco and the Wappner lab for discussions, and Andrés Rossi for technical support with confocal microscopy. This work was supported by Agencia Nacional de Promoción Científica y Tecnológica (ANPCyT) grants PICT-2015-0372 and PICT-2017-1356 to P.W.

## References

1. D. Voet, J. G. Voet, NewYork: John Wiley& SonsInc, 492 (2011).

2. J. G. Richards, in Fish physiology. (Elsevier, 2009), vol. 27, pp. 443–485.

3. M. Grieshaber, I. Hardewig, U. Kreutzer, H.-O. Pörtner, in Reviews of Physiology, Biochemistry and Pharmacology, Volume 125. (Springer, 1993), pp. 43–147.

4. L. Schito, S. Rey, Trends in cell biology 28, 128 (2018).

5. G. L. Semenza, The Journal of clinical investigation 123, 3664 (2013).

6. E. Klipp, B. Nordlander, R. Krüger, P. Gennemark, S. Hohmann, Nature biotechnology 23, 975 (2005).

7. H. Bisswanger, Perspectives in Science 1, 41 (2014).

8. E. Riekeberg, R. Powers, F1000Research 6, (2017).

9. N. Smolina, J. Bruton, A. Kostareva, T. Sejersen, Cell Viability Assays: Methods and Protocols, 79 (2017).

10. A. San Martín et al., PloS one 8, e57712 (2013).

11. A. San Martín et al., PloS one 9, e85780 (2014).

12. C. Zhang, Z.-H. Wei, B.-C. Ye, Applied microbiology and biotechnology 97, 8307 (2013).

13. P. Mächler et al., Cell metabolism 23, 94 (2016).

14. A. Gonzalez-Gutierrez, A. Ibacache, A. Esparza, L. F. Barros, J. Sierralta, bioRxiv, 610196 (2019).

15. K. Aoki, N. Komatsu, E. Hirata, Y. Kamioka, M. Matsuda, Cancer science 103, 614 (2012).

16. T. Zimmermann, in Microscopy Techniques. (Springer, 2005), pp. 245–265.

17. B. Phypers, Continuing education in Anaesthesia, critical care & pain 6, 129 (2006).

18. F. M. Zwiebel, U. Schwabe, M. S. Olson, R. Scholz, Biochemistry 21, 346 (1982).

19. M. Willenborg, U. Panten, I. Rustenbeck, European journal of pharmacology 607, 41 (2009).

20. M. Marron, T. Markow, K. Kain, A. Gibbs, Journal of Insect Physiology 49, 261 (2003).

21. V. Callier, S. C. Hand, J. B. Campbell, T. Biddulph, J. F. Harrison, Journal of Experimental Biology 218, 2927 (2015).

22. A. Casali, E. Batlle, Cell stem cell 4, 124 (2009).

23. N. S. Chandel et al., Journal of Biological Chemistry 275, 25130 (2000).

24. W. Ehleben et al., Respiration physiology 114, 25 (1998).

25. S. Rion, T. J. Kawecki, Journal of evolutionary biology 20, 1655 (2007).

26. A. G. Gibbs, L. A. Reynolds, in Comparative physiology of fasting, starvation, and food limitation. (Springer, 2012), pp. 37–51.

27. J. P. Monserrate, M. Y. Chen, C. B. Brachmann, BMC biology 10, 63 (2012).

28. M. Melani et al., Molecular biology of the cell 28, 3070 (2017).

29. J. Stelling, U. Sauer, Z. Szallasi, F. J. Doyle III, J. Doyle, Cell 118, 675 (2004).

30. J. E. Bailey, Nature biotechnology 17, 616 (1999).

31. O. Warburg, Science 123, 309 (1956).

32. S. N. Villegas, Disease Models & Mechanisms 12, dmm039032 (2019).

33. C.-W. Wang, A. Purkayastha, K. T. Jones, S. K. Thaker, U. Banerjee, Elife 5, e18126 (2016).

34. V. Slaninova et al., Open biology 6, 150155 (2016).

35. D. Bilder, M. Li, N. Perrimon, Science 289, 113 (2000).

36. D. Bilder, Genes & development 18, 1909 (2004).

37. T. Eichenlaub et al., Current Biology 28, 3220 (2018).

38. J. Aubin, Journal of Histochemistry & Cytochemistry 27, 36 (1979).

39. L. Szablewski, Biochimica et Biophysica Acta (BBA)-Reviews on Cancer 1835, 164 (2013).

40. H. R. Christofk et al., Nature 452, 230 (2008).

41. S. Desai et al., Oncotarget 5, 8202 (2014).

42. J. C. Schell et al., Molecular cell 56, 400 (2014).

43. D. K. Bricker et al., Science 337, 96 (2012).

44. G. Baek et al., Cell reports 9, 2233 (2014).

45. T. Hitosugi et al., Molecular cell 44, 864 (2011).

46. J. Kaplon et al., Nature 498, 109 (2013).

47. P. Miao, S. Sheng, X. Sun, J. Liu, G. Huang, IUBMB life 65, 904 (2013).

