## Supplementary Data for "A genetic toolkit for the analysis of metabolic changes in *Drosophila* provides new insights into metabolic responses to stress and malignant transformation"

#### Supplementary Information

##### Utilization of a linear unmixing algorithm to correct background and autofluorescence in FRET images

Autofluorescence is usually assumed to be a constant –and minor- component of the aquired fluorescence signal. However, this is not the case in *Drosophila* tissues that -due to their composition and complexity- show high and inhomogeneous autofluorescence (Sup Fig 1A). Autofluorescence is particularly problematic when assessing signal ratios, such as those characteristic of FRET experiments, especially when the intensity is low. In the current work, we estimated the autofluorescence in the donor and acceptor emission channels utilizing a spectral unmixing algorithm, followed by pixel-by-pixel substraction of the autofluorescence value obtained. The assumption in this method is that the fluorescence measured in a given channel (for example, YFP, $I_m^Y$) is the result of the sum of the fluorophore signal ($I_f^Y$) plus autofluorescence (in this example, detected in the YFP channel) ($I_{af}^Y$) (eq 1).

$$I_m^Y(x,y) = I_f^Y(x,y) + I_{af}^Y(x,y) \qquad (1)$$

Since direct determination of $I_{af}^Y$ is not possible due to its overlap with $I_f^Y$, $I_{af}^Y$ is estimated from the fluorescence intensity in a different emission wavelength, where no fluorophore signal can be detected. We call this third channel, defined by donor excitation wavelength and only autofluorescence emission, "A". $I_{af}^A$ can be then directly measured by taking a dedicated image in this particular channel (no additional illumination is necessary).

Estimation of the fluorescence intensity in the YFP channel ($I_{af}^Y$) from the autofluorescence signal in the AF channel ($I_{af}^A$) is possible due to the property of autofluorescence of having a wide emission spectrum that is reproducible for a given tissue. This happens because the source of autofluoresence is a complex mixture of different molecules (*38*), in contrast to a single fluorophore such as YFP. Thus the autofluorescence emission spectrum can be characterized in un-labeled samples, establishing a relation between the measured intensity in the YFP ($I_{af}^Y$) and the AF ($I_{af}^A$) channels that will be maintained in labeled samples. This method assumes that the relation between the fluorescence intensities detected in both channels is linear. Estimation of $I_{af}^Y$ is then performed by weighting the $I_{af}^A$ by a constant K (eq 2).

$$I_{af}^Y(\mathrm{x,y}) = \mathrm{K} * I_{af}^A(x,y) \qquad (2)$$

Noteworthy, while the fluorescence intensity from the fluorophore ($I_{af}^Y$), and autofluorescence ($I_{af}^A$) are functions of space, the constant K depends only on tissue properties and image settings. Then, eq.1 can be combined with eq 2, to get the correction equation (eq 3).

$$I_f^Y(x,y) = I_m^Y(x,y) - K * I_{af}^A(x,y) \qquad (3)$$

To apply this algorithm, then it is necessary to *i*) estimate K, and *ii*) acquire a specific image, $I_{af}^A$ in each experiment. For characterization of the autofluorescence emission spectrum and estimation of K, images from each of the three channels were obtained from tissues that do not express any sensor –where all the emission arises from autofluorescence- (Sup Fig 1B).

When working with sensor-expressing tissues, the autofluorescence was corrected according to eq. 3, and the ratiometric FRET signal (F) was calculated as in eq. 4

$$F(x,y) = \frac{I_f^Y(x,y)}{I_f^C(x,y)} \qquad (4)$$

By using eq 4, it is possible to calculate a FRET signal for each pixel, thereby generating the FRET map of the sample.

### Sup Fig 1

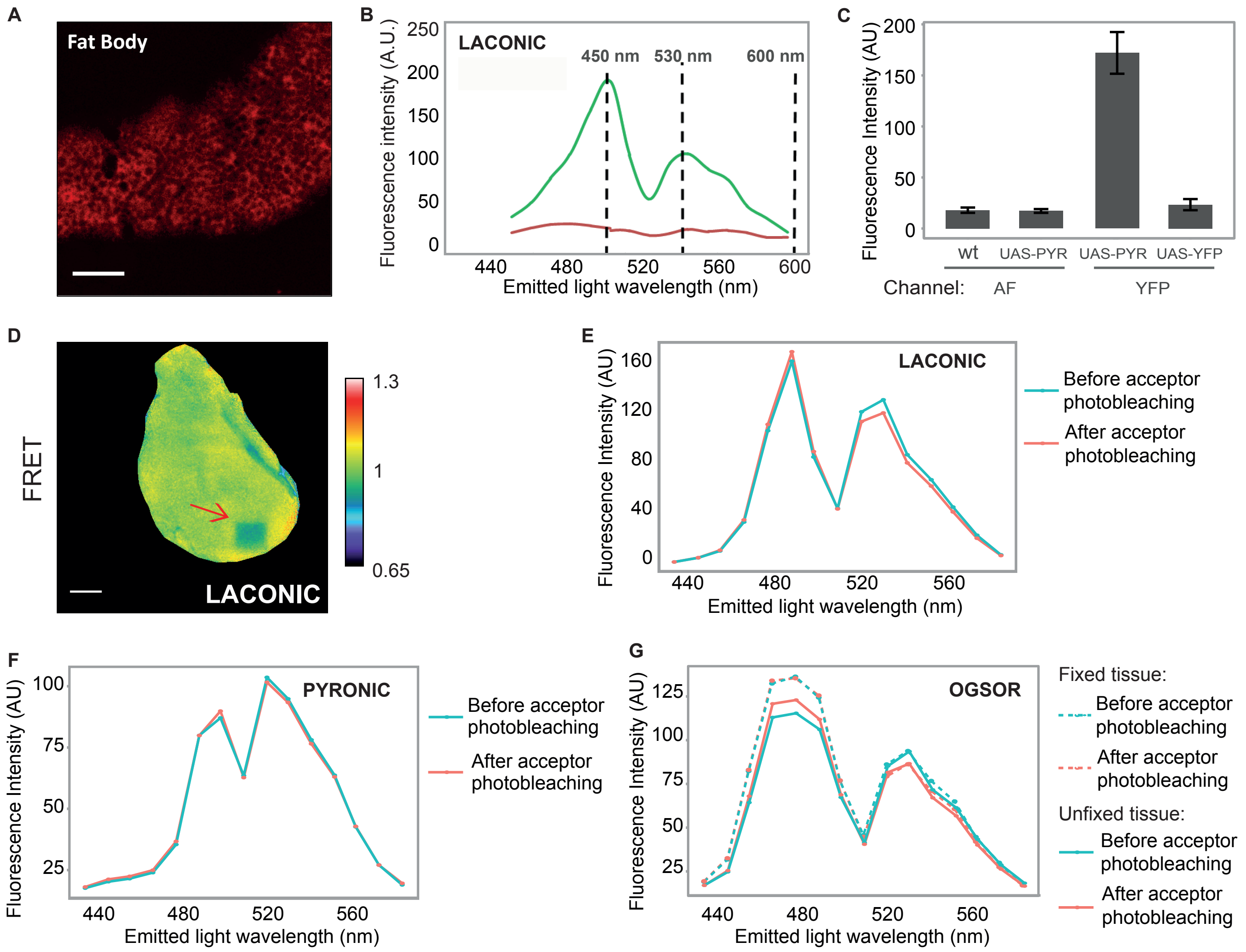

A
Fat Body
B
LACONIC
450 nm
530 nm
600 nm
Fluorescence intensity (A.U.)
Emitted light wavelength (nm)
C
Fluorescence Intensity (AU)
wt
UAS-PYR
UAS-PYR
UAS-YFP
Channel:
AF
YFP
D
FRET
1.3
1
0.65
LACONIC
E
LACONIC
Fluorescence Intensity (AU)
Emitted light wavelength (nm)
Before acceptor photobleaching
After acceptor photobleaching
F
PYRONIC
Fluorescence Intensity (AU)
Emitted light wavelength (nm)
Before acceptor photobleaching
After acceptor photobleaching
G
OGSOR
Fluorescence Intensity (AU)
Emitted light wavelength (nm)
Fixed tissue:
Before acceptor photobleaching
After acceptor photobleaching
Unfixed tissue:
Before acceptor photobleaching
After acceptor photobleaching

### Sup Fig 2

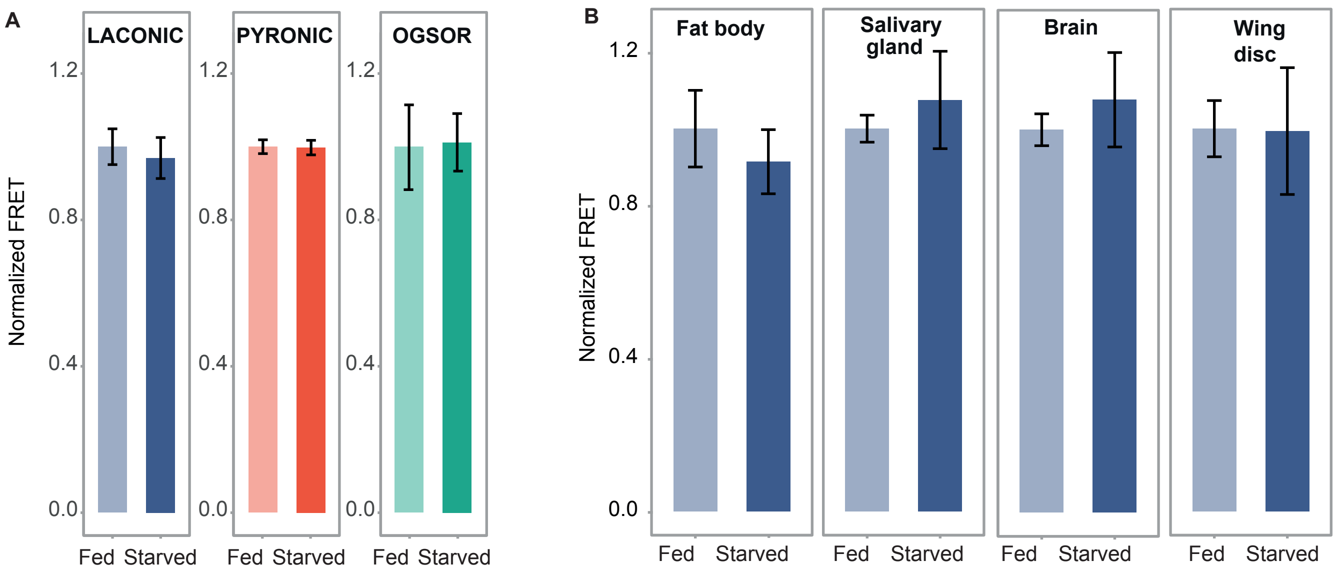
A
LACONIC
PYRONIC
OGSOR
Normalized FRET
1.2
0.8
0.4
0.0
Fed
Starved
B
Fat body
Salivary gland
Brain
Wing disc

### Sup Fig 3

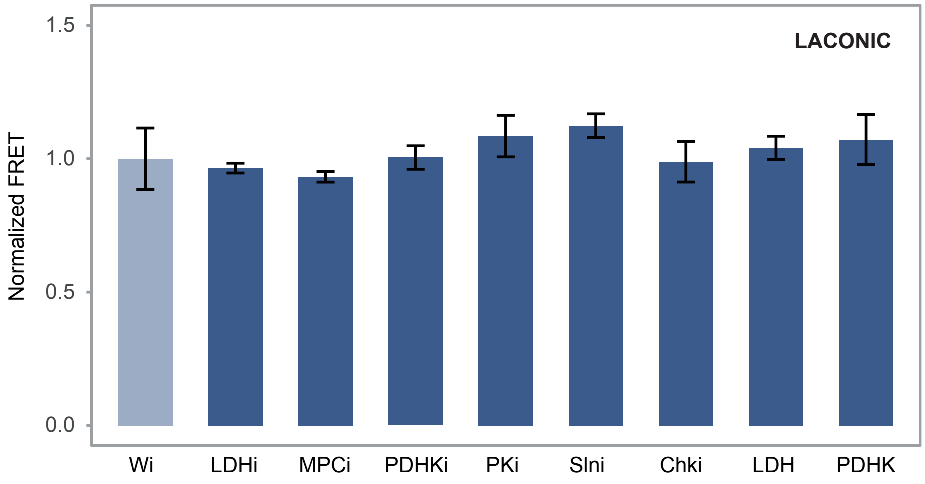

LACONIC
Normalized FRET
1.5
1.0
0.5
0.0
Wi
LDHi
MPCi
PDHKi
PKi
SIni
Chki
LDH
PDHK

**Supplementary Figure 1: Validation of FRET signals.** A) Autofluorescence at 600 nm of a wild type fat body upon excitation at 405 nm. Scale Bar: 50 μm. B) Emission spectra of wild type (red) or Laconic-expressing (green) wing discs upon excitation at 405 nm. The three channels relevant to the linear unmixing algorithm are highlighted with vertical dotted lines: CFP-donor channel (490 nm), YFP-acceptor channel (530 nm) and autofluorescence (AF) channel (600 nm). C) Fluorescence intensity at 600 nm of wild type wing discs (AF), at 600 nm of Pyronic-expressing discs (AF - Pyr), at 530 nm of Pyronic-expressing discs (YFP – Pyr) and at 530 nm of YFP-expressing discs (YFP), upon illumination at 458 nm. D) Laconic FRET map of a wing disc in which YFP has been photobleached by irradiating with high fluence at 488 nm in a small region of the posterior compartment (arrow). Scale bar: 50 μm. E) Emission spectra of Laconic-expressing discs upon excitation at 458 nm before or after YFP photobleaching. F) Emission spectra of Pyronic-expressing discs upon excitation at 458 nm before or after YFP photobleaching. G) Emission spectra of OGsor-expressing discs upon excitation at 458 nm before or after YFP photobleaching, either in fixed (dotted lines) or unfixed samples (solid lines).

**Supplementary Figure 2: Metabolite levels remain constant in wing discs when larvae are subjected to nutrient deprivation.** A) Laconic, Pyronic and OGsor FRET signals from wing imaginal discs of $3^{rd}$ instar larvae fed *ad libitum* or subjected to 6 h nutrient deprivation. No differences in lactate levels can be detected. B) Laconic FRET signal from $3^{rd}$ instar wing discs, brain, salivary glands and fat body fed *ad libitum* or subjected to a 18 h nutrient deprivation. No differences in lactate levels can be detected in this case either. Data represent the media +/- SD. n≥20 per group.

**Supplementary Figure 3: Single genetic manipulations do not alter lactate levels.** Laconic FRET signal of wing discs where the indicated proteins have been either overexpressed or silenced by expression of specific RNAis with an En-Gal4 driver. Note that these single genetic manipulations are not sufficient to alter lactate levels. Data represent the media +/- SD. n≥20 per group.
